# *Toxoplasma* type II effector GRA15 has limited influence *in vivo*

**DOI:** 10.1101/2024.02.23.581829

**Authors:** Emily F. Merritt, Joshua A. Kochanowsky, Perrine Hervé, Alison A. Watson, Anita A. Koshy

## Abstract

*Toxoplasma gondii* is an intracellular parasite that establishes a long-term infection in the brain of many warm-blooded hosts, including humans and rodents. Like all obligate intracellular microbes, *Toxoplasma* uses many effector proteins to manipulate the host cell to ensure parasite survival. While some of these effector proteins are universal to all *Toxoplasma* strains, some are polymorphic between *Toxoplasma* strains. One such polymorphic effector is GRA15. The *gra15* allele carried by type II strains activates host NF-κB signaling, leading to the release of cytokines such as IL-12, TNF, and IL-1β from immune cells infected with type II parasites. Prior work also suggested that GRA15 promotes early host control of parasites *in vivo*, but the effect of GRA15 on parasite persistence in the brain and the peripheral immune response has not been well defined. For this reason, we sought to address this gap by generating a new IIΔ*gra15* strain and comparing outcomes at 3 weeks post infection between WT and IIΔ*gra15* infected mice. We found that the brain parasite burden and the number of macrophages/microglia and T cells in the brain did not differ between WT and IIΔ*gra15* infected mice. In addition, while IIΔ*gra15* infected mice had a lower number and frequency of splenic M1-like macrophages and frequency of PD-1+ CTLA-4+ CD4+ T cells and NK cells compared to WT infected mice, the IFN-γ+ CD4 and CD8 T cell populations were equivalent. In summary, our results suggest that *in vivo* GRA15 may have a subtle effect on the peripheral immune response, but this effect is not strong enough to alter brain parasite burden or parenchymal immune cell number at 3 weeks post infection.

## Introduction

To successfully establish a persistent infection, a microbe must take a “Goldilocks” route. The microbe must evade host defenses enough to avoid microbial elimination while also preventing host death from an overwhelming microbial burden or immune response. Thus, successful persistent microbes evolve mechanisms for provoking “the right” amount of a host response (1,2). *Toxoplasma gondii* is an eukaryotic intracellular parasite that persistently infects many warm blooded animals—from birds to humans—including approximately 10-15% of the United States population (3). *Toxoplasma* has achieved such success, in part, by manipulating host cell signaling pathways through a variety of secreted effector proteins. These secreted effector proteins are often known as ROPs and GRAs and are delivered by specialized secretory organelles. Different ROPs and GRAs directly block immune clearance, alter the host cell cycle, drive cytoskeletal remodeling, and alter apoptotic pathways (4–11). While many of these effector proteins are the same in all *Toxoplasma* strains, some are polymorphic and show *Toxoplasma* strain-specific effects (12,13). One such polymorphic effector protein is GRA15 (14).

During *in vitro* infection with type II *Toxoplasma* strains—but not with type I or type III strains—GRA15 activates the NF-κB pathway, which leads to IL-12, IL-1β, and TNF release by macrophages (15–18). GRA15 also limits parasite growth in IFN-γ stimulated human and murine fibroblasts *in vitro* by recruiting host defense proteins to the parasite’s intracellular niche (15). Consistent with GRA15 stimulating pro-inflammatory host responses that limit parasite expansion, during acute infection, mice inoculated with type II parasites that lack GRA15 have lower local IFN-γ levels and higher parasite burdens compared to mice infected with wild-type type II parasites (14).

While such findings might be expected to result in a higher systemic and brain parasite burden during later stages of infection, the data are mixed. One paper found that IIΔ*gra15* parasites showed no difference in cyst counts at 21 days post infection (dpi) compared to parental parasites, while another paper found that IIΔ*gra15* parasites showed a trend toward a decrease in cyst count compared to WT parasites (19,20). Given these discrepant studies, we sought to re-address the role of GRA15 in outcomes of type II infection, including assessing the systemic and brain immune response as well as the brain parasite burden.

## Methods

### Ethics Statement

All procedures and experiments were carried out in accordance with the Public Health Service Policy on Human Care and Use of Laboratory Animals and approved by the University of Arizona’s Institutional Animal Care and Use Committee (#12–391). All mice were bred and housed in specific-pathogen-free University of Arizona Animal Care facilities.

### Parasite maintenance and generation of IIΔ*gra15* and IIΔ*gra15*::GRA15

All parasite strains were maintained through serial passage in human foreskin fibroblasts (HFFs) in DMEM supplemented with 10% FBS, 100 I.U./ml penicillin/streptomycin, and 2 mM glutagro. All parasite strains were generated from a type II strain Pruginaud (PruΔ*hpt*) in which the endogenous hypoxanthine xanthine guanine phosphoribosyl transferase gene is deleted. The wild type (WT) strain used throughout the paper expresses a Cre fusion protein that is injected into host cells prior to parasite invasion (21). The IIΔ*gra15* used throughout this work also expresses the Cre fusion protein.

To disrupt GRA15 in IIΔ*hpt* parasites, GRA15 targeting CRISPR plasmids (sgGRA15Up and sgGRA15down) were generated from a sgUPRT plasmid (plasmid #54464) using a Q5 mutagenesis protocol (22). To generate a plasmid to insert *hpt* and *toxofilin:cre* into the GRA15 locus, upstream (500-bp) and downstream (500-bp) adjacent to the sgGRA15Up and sgGRA15Down target sequences were used to flank *hpt* and *toxofilin:cre*. We then transfected the IIΔ*hpt* parasites with the 1) sgGRA15Up CRISPR, 2) sgGRA15Down CRISPR, and 3)pTKO plasmid (23) with GRA15 homology regions flanking *hpt* and *toxofilin:cre*. These parasites underwent selection using media containing 25 mg/ml mycophenolic acid and 50 mg/ml of xanthine prior to dilution to individual clones (24). Single clones were then screened for disruption of the *gra15* locus and confirmed to have lost NF-κB activation by immunofluorescence. Clones were also confirmed to express *toxofilin:cre* by causing Cre-mediated recombination as previously described (25).

A complemented IIΔ*gra15*::GRA15 strain was made by inserting the GRA15 coding sequence with 1000 bp upstream of the GRA15 TSS into a plasmid containing the selectable marker bleomycin (26). The plasmid was linearized and transfected into IIΔ*gra15* parasites. These parasites were placed under selection in complete DMEM supplemented with 5 µg/ml zeocin until lysing out. Lysed out parasites were incubated in 50 µg/ml zeocin media for 4 hours before being transferred to HFFs containing the 5 µg/ml zeocin media. This process was repeated three times prior to cloning by limiting dilution. Single clones were then screened for expression of *gra15* by Q-PCR and the ability to activate NF-κB pathway by immunofluorescence.

### Mice

Unless specifically noted, mice used in this study are Cre-reporter mice in a C57Bl/6J background. Cells in these mice express GFP when cells undergo Cre-mediated recombination (27). These mice were purchased from Jackson labs and bred in the University of Arizona Animal Center (stock # 007906). BALB/cJ mice (Strain #:000651) were used for one experiment. Male and female mice were intraperitoneally inoculated with 10,000 syringed lysed parasites resuspended in 200 µl of UPS grade PBS. Unless otherwise stated, two cohorts were used for each experiment. For 3 week post infection (wpi) studies, cohort one included 4-5 mice per infection, aged 12-16 weeks, with initial weights between 18 and 33 grams. Cohort two included 9-12 mice per infection, aged 6-10 weeks, with initial weights between 16 and 32 grams. For acute time points of 2- and 5-days post infection, each cohort contained 4-5 mice per infection. Mice were given food and water ad libitum and provided moist chow to alleviate suffering.

### Tissue preparation for histology and DNA extraction

At the appropriate time points, mice were euthanized with CO_2_, without use of anesthesia, and transcardially perfused with 20 ml cold PBS. Brains were removed and divided into two hemispheres. The left hemisphere was drop fixed by placement in 4% paraformaldehyde (PFA). The next day, PFA was removed and replaced with 30% sucrose. After sucrose embedding, brains were sagittally sectioned to 40 µm sections using a microtome (Microm HM 430) and stored in cryoprotectant media at -20° C until staining. The anterior ¼ of the right half of the brain was sectioned coronally, placed in an Eppendorf tube, and flash frozen until used for DNA extraction.

### NF-κB activation assay

Syringe lysed parasites were added to confluent HFF monolayers grown on glass coverslips at an MOI of 7.5 and spun down at 300 rpm for 1 minute. 24 hours post infection, cells were washed and fixed for 15 minutes with 4% PFA followed by 5 min in ice cold methanol. Cells were then blocked in 3% goat serum for 1 hour at room temperature and incubated in mouse anti-Sag1 (28) [DG52] (gift from John Boothroyd, 1:5000) and anti-NF-κB (p65) (Santa Cruz Biotechnology, sc-372, 1:1000) antibodies overnight at 4°C. The next day, cells were washed to remove excess antibody and incubated in goat anti-mouse secondary antibody AF568 (Thermo Fischer Scientific, A-11004, 1:500) and goat anti-rabbit AF488 (Life technologies, A-11008, 1:500) for one hour. Coverslips were then washed 3 times in PBS, with the first wash containing Hoechst (1:5000) to stain for host cell and parasite nuclei. Images were then obtained on an ECHO Revolve fluorescent microscope to analyze nuclear NF-κB localization.

To measure NF-κB activation at early time points, syringe lysed parasites were filtered and washed in 40 ml of cDMEM prior to addition to confluent HFF monolayers grown on glass coverslips at an MOI of 5. Cells were fixed in 4% PFA at 1, 3, or 24 hrs post infection, blocked in 3% goat serum for 1 hour at room temperature and incubated in mouse anti-Sag1(28) [DG52] and anti-NF-κB (Cell Signaling Technology, 8242S, 1:1000 antibodies) overnight at 4°C. The subsequent steps followed the protocol described above.

### Growth assay

Syringe lysed parasites were added to confluent HFF monolayers grown on glass coverslips at an MOI of 1 and spun down at 300 rpm for 1 minute. 24 hours post infection, cells were washed in PBS and fixed for 20 minutes in 4% PFA. Cells were then permeabilized, blocked, and stained using an anti-*Toxoplasma* antibody (Thermo Fischer, PA17252, 1:5000, goat anti-rabbit 568 Thermo Fischer, A11011, 1:500). To enumerate the number of parasites per vacuole, coverslips were analyzed using an ECHO Revolve fluorescent microscope.

### Plaque assay

Confluent monolayers of HFF cells were infected with 250 parasites of the indicated strains in cDMEM. After 10 days, media was removed, cultures were washed with PBS, and the monolayers were fixed in ice cold methanol for 10 minutes. Fixed monolayers were then stained with crystal violet for 10 minutes at room temperature.

### Immunohistochemistry

For identification of macrophages/microglia, free floating brain sections were washed, treated in H_2_O_2_ for 40 minutes, washed again, blocked with goat serum, and incubated with polyclonal rabbit anti-Iba-1 antibody overnight (Wako Pure Chemical Industries, 019-19741 Ltd. (1:3000)). The next day, samples were washed, incubated for 1 hour in biotinylated goat anti-rabbit antibody (Vector Laboratories, BA-1000 (1:500)). After washing off residual secondary antibody, samples were incubated in ABC solution (Thermo Fisher, 32020) for 1 hour followed by 60 second treatment with 3,3’-Diaminobenzidine (DAB)(Vector Laboratories, SK-4100). Samples were then washed, mounted, and cover slipped prior to immune cell quantification.

### Immunofluorescence

For identification of T cells, free floating brain sections were washed in TBS, blocked with 3% goat serum diluted in TBS for 1 hour, and then incubated overnight with hamster anti-CD3ε antibody diluted in 1% goat serum/0.3% Triton-X100/TBS (BD Biosciences, 550277). The next day samples were washed in TBS and incubated at room temperature in goat anti-hamster 647 (Life Technologies, A-21451). After a 4-hour incubation in secondary antibody, samples were washed for 5 minutes in TBS/Hoechst (1:5000) followed by 2 subsequent washes in TBS. Brain sections were then mounted, cover slipped with Fluoromount-G™ (Southern Biotech, 0100-01), and z-stacks were obtained on an ECHO Revolve microscope using a 10x objective.

### Iba-1+ cell quantification

To quantify Iba-1+ cells in brain sections, stained sections were imaged using light microscopy. Eight images were obtained in a stereotyped pattern within the cortex of the brain section using a 20x objective. Three matched sections were imaged per mouse (24 images/mouse). Cells were quantified manually using FIJI software. Individuals quantifying cells were blinded to infection status of mice.

### T cell quantification

Imaris software was used to quantify number of T cells within each 40 µm confocal image. The spots tool was used to generate a threshold of detectable T cells and quantified by the program. Individuals quantifying cells were blinded to infection status of mice.

### Quantitative PCR

To quantify parasite burden, genomic DNA was isolated from the anterior quarter of the right hemisphere (brain), the left lobe of the liver, or the distal quarter of the spleen using DNeasy Blood and Tissue kit (Qiagen, 69504), following the manufacturer’s protocol. The *Toxoplasma* B1 gene was amplified using SYBR Green on the Eppendorf Mastercycler et realplex 2.2 system. Gapdh was used to normalize parasite DNA levels.

### Cyst Stain

Sagittal brain sections were blocked in 3% goat serum diluted in 0.3% TritonX-100/TBS for 1 hour. These sections were then incubated with biotinylated Dolichos Biflorus Agglutinin (DBA) (Vector laboratories 1031, 1:500) and a polyclonal rabbit anti-*Toxoplasma* antibody (Thermo Fisher Scientific, PA17252, 1:5000) overnight at 4° C. Samples were then washed and incubated with Streptavidin Cy5 (Life technologies, S21374, 1:500) and goat anti-rabbit 568 secondary (Thermo Fisher Scientific, A11011, 1:500) for 4 hours at room temperature, after which samples were washed to remove residual antibody. Hoechst (Thermo Fisher Scientific, H3570, 1:5000) was added to the first TBS wash for 5 minutes to stain for nuclei. Sections were then washed two more times, mounted on slides, and cover slipped using Fluoromount-G™. The number of cysts (DBA+, anti-*Toxoplasma* antibody+) was enumerated using an Echo Revolve fluorescent microscope.

### Single cell suspension for Flow Cytometry

At appropriate time points, mice were euthanized by CO_2_ and intracardially perfused with 20 ml cold PBS. Spleens were then harvested for flow cytometry, maintained in complete RPMI (86% RPMI, 10% FBS, 1% penicillin/streptomycin, 1% L-glutamine, 1% NEAA, 1% sodium pyruvate, and <0.01% β-mercaptoethanol) and processed to generate single cell suspensions. For single cell suspension, spleens were passed through a 40 µm strainer and centrifuged at 1200 rpm, 4°C, for 5 minutes. After removal of supernatant, red blood cells were lysed by addition of 1 ml ammonium chloride-potassium carbonate (ACK) lysis buffer (Life Technologies, A1049201). ACK was neutralized by the addition of cRPMI, centrifuged at 1200 rpm, 4°C, for 5 minutes. The supernatant was removed and the pellet resuspended in cRPMI. The number of viable cells was quantified by diluting 10 µl of the single cell suspension in 90 µl trypan blue and counting on a hemocytometer. T cell panels to be quantified for IFN-γ were treated with PMA, Ionomycin, and Brefeldin for 4 hours in 37°C incubator prior to washing, blocking, and staining.

### Staining for Flow Cytometry

One million live cells of each sample were plated into a 96 well plate, washed in FACS buffer (1% FBS/PBS), and blocked with Fc block (Biolegend, 101302) to prevent nonspecific staining. Samples were then stained for a T cell panel or a macrophage panel. Samples were incubated in antibody (1:100) diluted in FACS buffer for at least 30 minutes, protected from light, then stained with live/dead Fixable yellow Dead Cell stain (Life Technologies, L34959). Samples were then washed and fixed using intracellular staining permeabilization and fixation kit (eBioscience, 00-5223-00). To stain for Foxp3, T-bet, Gata-3, and IFN-γ, the manufacturer’s intracellular staining protocol was used (eBioscience, 00-5223-56; 00-5123-43; 00-8333-56). Samples were then washed, run on a LSRII (University of Arizona Cancer Center Flow Cytometry Core), and data analyzed using FlowJo™ Software.

### Peritoneal Exudate Cells Isolation

Cre reporter mice were inoculated intraperitoneally with saline or 10,000 WT or II*Δgra15* parasites. At 2 and 5 dpi, peritoneal exudate cells were collected by injecting 5 ml of cold PBS into the exposed peritoneal cavity, massaging the cavity, and recollecting PBS/cellular suspension. PECS were then incubated in Fc block, stained for CD45, and run on the LSRII.

### Parasite RNA isolation

Confluent human foreskin fibroblasts were infected with indicated strains for 48 hours. Monolayers were scraped, syringe lysed, and resuspended in TRIzol™; RNA was extracted per manufacturer’s instructions (Thermo Fisher Scientific, 15596026). One µg of isolated RNA was converted to cDNA using High-Capacity cDNA Reverse Transcriptase kit (Thermo Fisher Scientific, 4368814). Q-PCR was performed on cDNA using GRA15 and TgActin specific primers.

### Statistics

Graphs were generated and statistical tests were run using Prism software version 9.4.1. All *in vivo* experiments in C57BL/6 mice were repeated with two independent cohorts; unless otherwise noted the data were analyzed with a two-way analysis of variance (ANOVA) with uncorrected Fisher’s LSD. Infection of BALB/c mice was done once; data were analyzed with a T-test. For intracellular growth assays and plaque assays, experiments were repeated three times and statistical analysis was conducted on the composite data. For intracellular growth assay, a two-way ANOVA with uncorrected Fisher’s LSD was used, for plaque assays a one-way ANOVA was used. Analysis of parasite genomes in the liver at 5 dpi showed one mouse from one cohort to be an outlier as determined by the ROUT outlier test. Therefore, that mouse was removed for statistical analysis.

## Results

### GRA15 does not influence parasite burden or macrophage/microglia and T cell abundance in the brain at 3 wpi

To probe the influence of GRA15 during early chronic infection, we generated a type II strain (Prugniaud or Pru) that lacked *gra15* (IIΔ*gra15*) and the appropriate complemented strain (IIΔ*gra15*::GRA15) using previously described CRISPR-Cas9 methodology (29). As the IIΔ*gra15* and IIΔ*gra15*::GRA15 strains express a rhoptry::Cre recombinase fusion protein, for the wild-type (WT)/control strain, we used a Pru strain that has been engineered to express the same rhoptry::Cre fusion protein (21). The IIΔ*gra15* was confirmed to lack NF-κB activation and the complemented strain restored NF-κB activity at 24 hours post infection (hpi) (**S1 Fig A, B**). At 1 and 3 hpi, none of the strains induced NF-κB nuclear localization, regardless of GRA15 expression (**S1 Fig C**). To assess GRA15’s effect *in vivo*, Cre reporter mice that express GFP only after Cre-mediated recombination (27) were inoculated with saline or WT, IIΔ*gra15*, or IIΔ*gra15*::GRA15 parasites. At 3 weeks post infection (wpi), spleen and brain were harvested. To assess overall brain parasite burden, we performed Q-PCR for a *Toxoplasma* specific gene (B1) on genomic DNA isolated from the brain (23,30). We found no statistical difference between WT and IIΔ*gra15* infected brain though the IIΔ*gra15*::GRA15 infected brain consistently showed a lower parasite burden (**Fig 1A**). As a second mechanism for assessing brain parasite burden, we quantified the number of cysts by staining brain sections with Dolichos biflorous agglutinin (DBA), a lectin that stains sugar moieties on components of the cyst wall (**Fig 1B**) (31). Consistent with the Q-PCR data, cyst counts from WT and IIΔ*gra15* infected brain were not statistically different while cysts counts from IIΔ*gra15*::GRA15 infected brain were lower (**Fig 1C**). Given that the IIΔ*gra15*::GRA15 strain consistently appeared to be less capable of establishing an *in vivo* infection in multiple cohorts of mice, we performed *in vitro* studies to determine if this strain had a growth defect and/or had an unusual expression of *gra15*. Indeed, the IIΔ*gra15*::GRA15 strain showed a replication defect at 24 hours post infection (**S1 Fig D)**, though this difference did not translate into a defect in plaque formation (**S1 Fig E-G**). In addition, we determined that the complemented strain expressed approximately 5 fold more *gra15* compared to the WT strain (**S1 Fig H**). Given the lytic cycle defect—which we expect would be exacerbated *in vivo*—and the increased expression of *gra15* in the IIΔ*gra15*::GRA15 strain, we decided to move forward without the complement, as these phenotypes introduce variables for which we cannot control.

**Figure 1.**
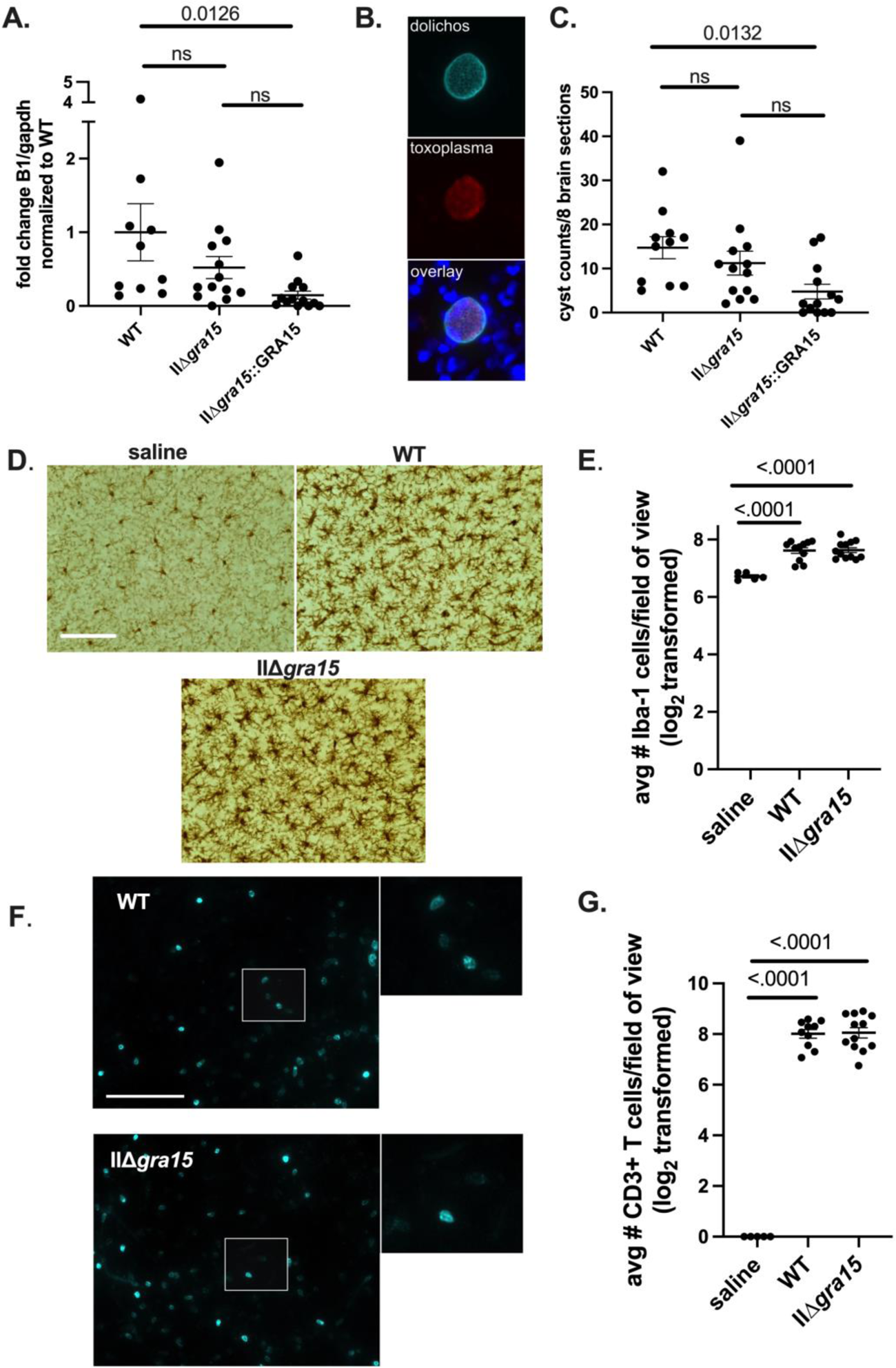
GRA15 does not influence brain parasite burden in the brain at 3 weeks post infection. Mice were intraperitoneally (i.p.) inoculated with saline (control) or 10,000 WT, IIΔ*gra15*, or IIΔ*gra15*:GRA15 parasites. Brains and spleens were harvested at 3 weeks post infection (wpi). Mice from these infections were used in Figures 1-5. **A.** Graph of *Toxoplasma* brain burden as assessed by Q-PCR for the *Toxoplasma*-specific B1 gene. **B.** Representative images of a brain tissue cyst stained with Dolichos biflorus agglutinin (DBA). *Top image* is DBA staining, *middle image* is staining with anti-*Toxoplasma* antibodies, and *bottom image* is merge. **C.** Quantification of cyst numbers in 8 brain sections per mouse. **D.** Representative images of Iba1+ cells (microglia/macrophages). Scale bar, 100 µm **E**. Quantification of the number of Iba-1+ cells. **F**. Representative images of CD3ε+ cells (T cells). Scale bar, 100 µm. Panels on right are enlarged insert of white box in left panels. **G**. Quantification of CD3ε+ cells. **A, C, E, G**. Bars, mean ± SEM. N = 8 fields of view/section, 3 sections/mouse, 5-12 mice/group. For each mouse, the number of cells/section was averaged to create a single point. Data representative of two independent experiments.

As GRA15 influences macrophage phenotypes *in vitro* and a change in macrophage skewing might affect the neuroinflammatory response without altering brain parasite burden, we next sought to evaluate the brain immune response. We focused on macrophages/microglia and T cells because these are the primary immune cells to infiltrate and/or be activated in the brain upon *Toxoplasma* infection (29). To quantify the number of macrophages/microglia, we stained tissue sections with anti-Iba1 antibodies, which stains a cytoskeletal protein (Iba1) expressed by both macrophages and microglia. We then quantified the number of Iba1+ cells manually (29) finding no difference in the number of Iba1+ cells in brain sections from WT and IIΔ*gra15* infected mice (**Fig 1D,E**). To quantify T cells, we performed immunofluorescent assays for T cells using an anti-CD3ε antibody. We then imaged the stained tissue sections and analyzed the images with Imaris software, which is capable of segregating and counting the stained T cells in an automated manner (**Fig 1F,G**). We found no difference in the number of CD3ε+ cells in brain sections from WT and IIΔ*gra15* infected mice. Collectively, these data suggest that GRA15 does not affect *Toxoplasma*’s dissemination to or persistence in the brain at 3 wpi. GRA15 also does not appear to alter the number of macrophage/microglia or T cells present in the brain at 3 wpi.

### GRA15 may influence M1-like polarization of macrophages at 3 wpi

While IHC allows us to quantify infiltrating immune cells, it cannot assess the polarization state of immune cells, which can be done by flow cytometry. Given that GRA15 induces an M1-like phenotype in infected macrophages *in vitro* (32), we were interested in determining how this gene influences macrophage phenotypes *in vivo*. As prior data from our lab has shown that the immune response within the spleen mirrors the immune response found in the brain at 3 wpi (29), we used splenocytes for our analyses. To that end, we used the following markers to segregate macrophages into pro-inflammatory macrophages (M1-like): CD45+, F4/80+/ CD11b^hi^ CD11c^lo/int^/ CD80+ CD86+ and wound-healing macrophages (M2): CD45+, F4/80+/ CD11b^hi^ CD11c^lo/int^/ CD206+/F4/80+ (gating scheme shown in **S2 Fig**). Given that we did not use iNOS staining which is required to identify a true M1 macrophage, we refer to our CD80+ CD86+ population as M1-like macrophages. Consistent with the *in vitro* data, our analyses showed that, compared to WT infected mice, IIΔ*gra15* infected mice have fewer M1 macrophages (**Fig 2A,B**). We did not detect differences in M2 macrophages (**Fig 2C,D**).

**Figure 2.**
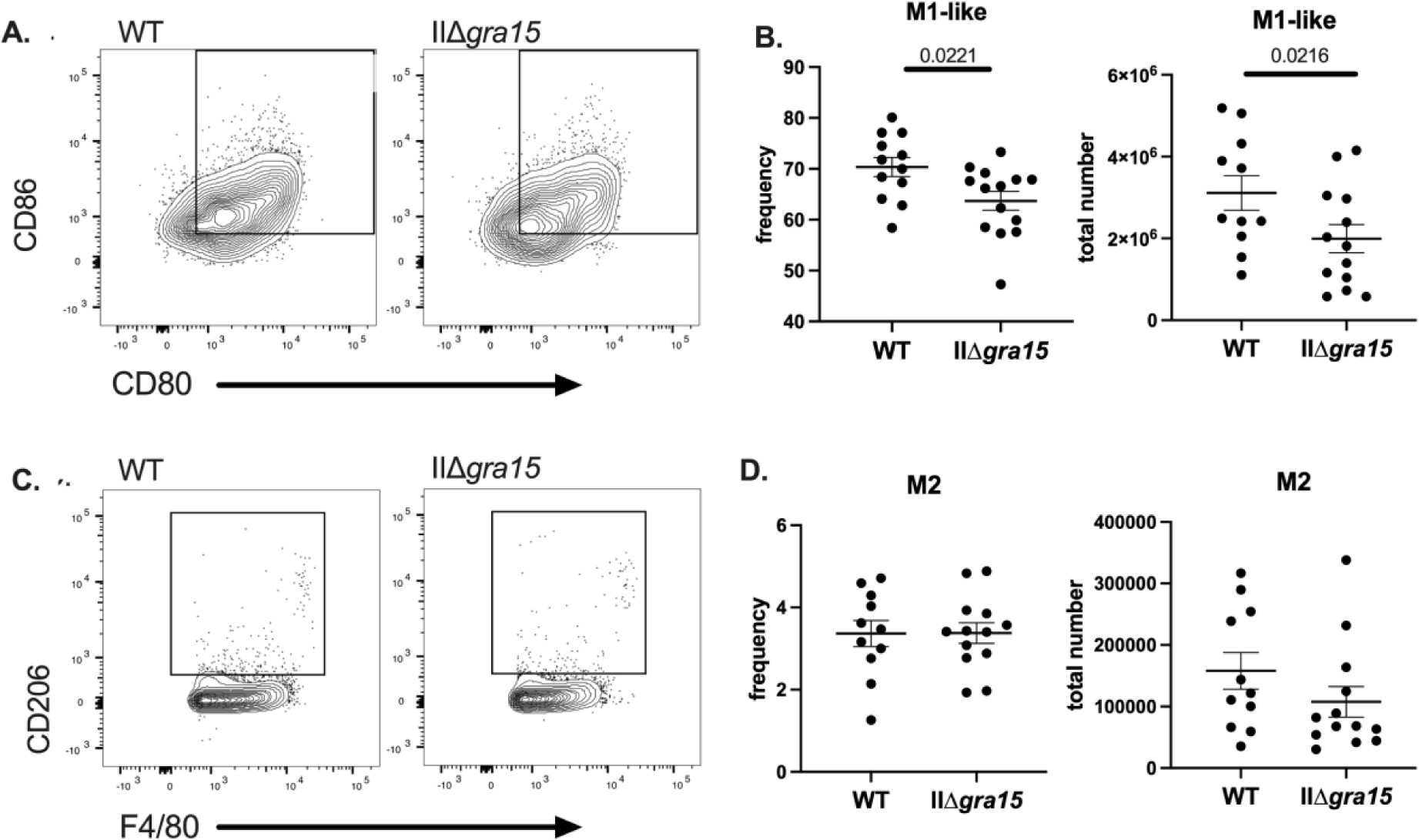
GRA15 may affect splenic M1-like macrophage population at 3 wpi. **A, B**. Splenic mononuclear cells were evaluated for the presence of M1-like macrophages (CD45+, F4/80+, CD11b^hi^, CD11b^lo/int^, CD80+, CD86+) **C, D**. Splenic mononuclear cells were evaluated for the presence of M2 macrophages (CD45+, F4/80+, CD11b^hi^, CD11c^lo/int^, CD206+ (MMR). Bars, mean ± SEM. N=11-12 mice/infected group. Data are representative of two independent experiments.

### GRA15 does not affect IFN-γ producing T cell populations at 3 wpi

M1/M1-like macrophages are expected to produce IL-12 (18,29,32). As IL-12 is one of many signals that polarizes naïve CD4+ T cells to be T-bet+, IFN-γ producing Th1 cells, we hypothesized that the lower number of M1-like macrophages provoked by IIΔ*gra15* parasites might result in decreases in IFN-γ production by T cells (33,34). To test this possibility, we profiled the splenic T cell compartment, assessing CD4 and CD8 numbers as well as their capabilities to produce IFN-γ (gating scheme is shown in **S3 Fig**). We found no differences between the groups in terms of the number or frequency of Th1, Th2, or Treg T cells (**Fig 3A-F**). The number of IFN-γ producing CD4 and CD8 T cells was also not different (**Fig 4**). Collectively, these data suggest that, at 3 wpi, GRA15 does not influence IFN-γ production in CD4 or CD8 T cells, despite potentially influencing M1-like macrophage number and frequency.

**Figure 3.**
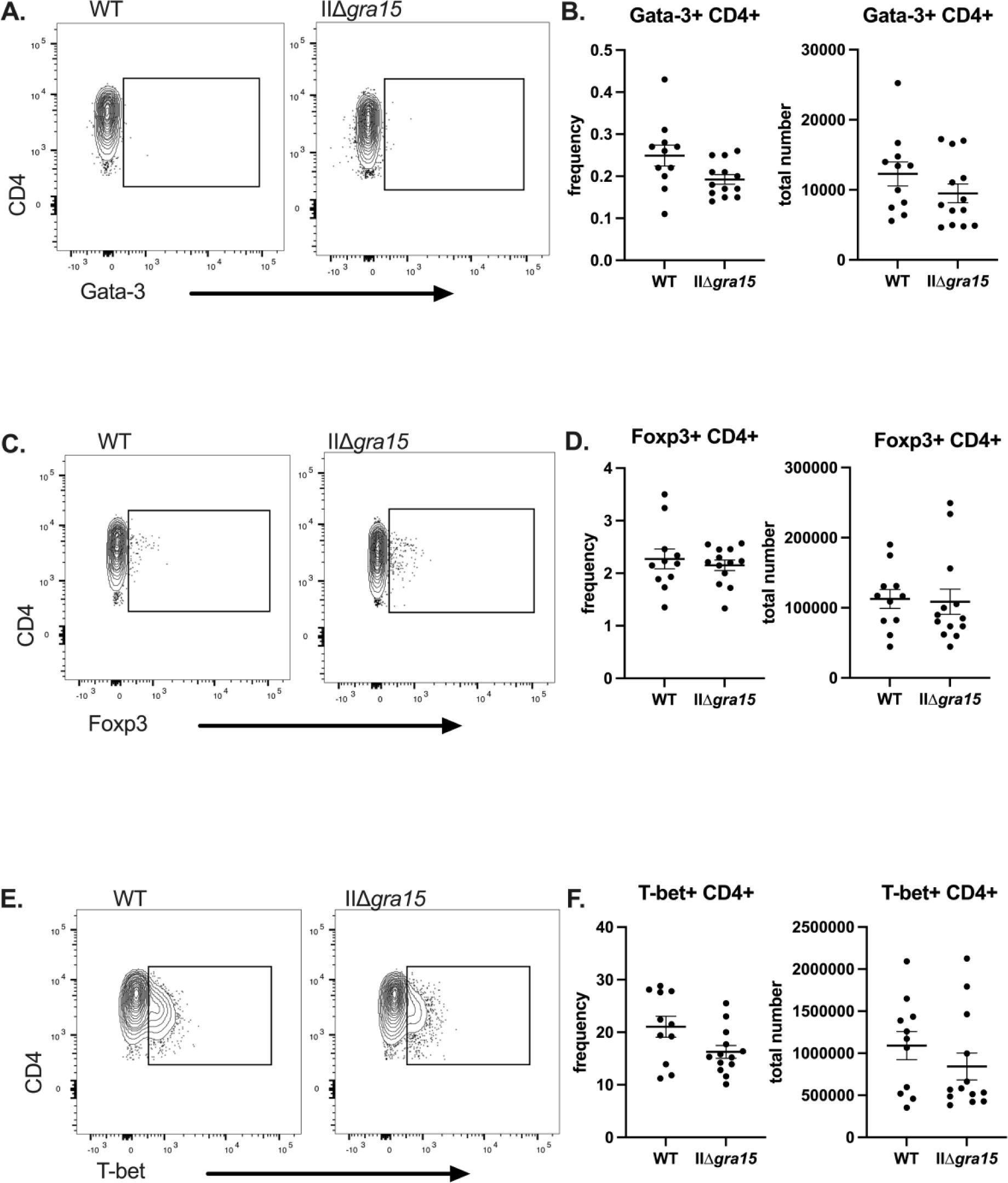
GRA15 does not influence splenic Th2, Tregs, or Th1 CD4+ T cells populations at 3 wpi. **A,B**. Splenic CD4+ CD3+ T cells were evaluated for the presence Th2 T cells (CD3+ CD4+ Gata-3+) **C,D.** Splenic CD4+ CD3+ T cells were evaluated for the presence regulatory T cells (CD3+ CD4+ Foxp3+) **E,F.** Splenic CD4+ CD3+ T cells were evaluated for the presence of Th1 T cells (CD3+ CD4+ T-bet+) Bars, mean ± SEM. N=11-12 mice/infected group. Unlisted p values were not significant. Data are representative of two independent experiments.

**Figure 4.**
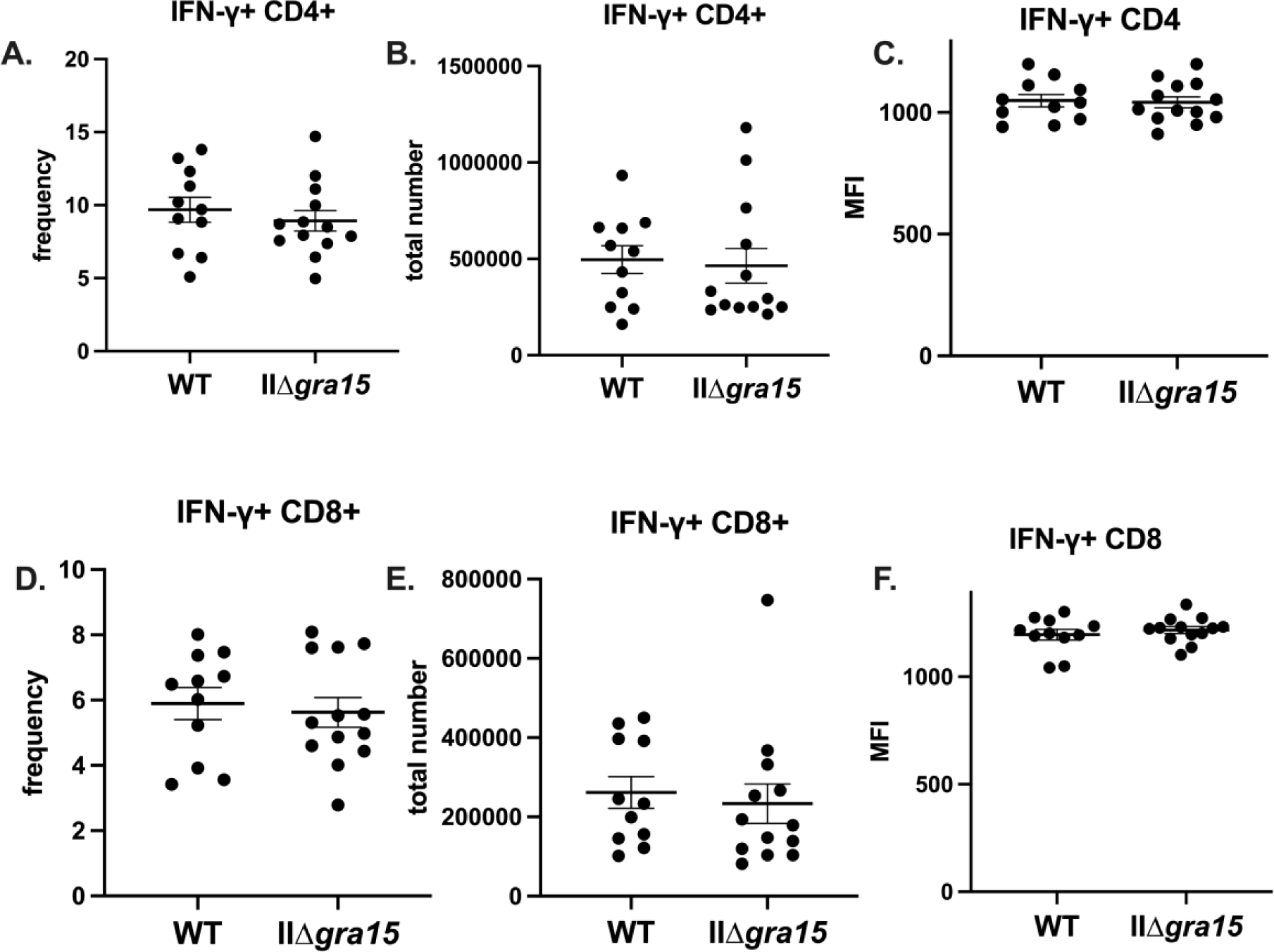
GRA15 does not influence IFN-γ producing T cell populations at 3 wpi. **A**. Splenic CD4+ CD3+ T cells were evaluated for their ability to make IFN-γ **B**. Splenic CD8+ CD3+ T cells were evaluated for their ability to make IFN-γ. Bars, mean ± SEM. N=11-12 mice/infected group. Data are representative of two independent experiments. Unlisted p values were ≥ 0.05.

### GRA15 may influence the frequency of peripheral “exhausted” T cells and NK cells at 3 weeks post infection in C57BL/6 mice

As work from other labs have identified T cell exhaustion during chronic time points of *Toxoplasma* infection (35,36) and because such analysis has not been done with Δ*gra15* strains, we assessed the T cell compartment for exhausted T cells by looking for co-expression of inhibitory markers PD-1 and CTLA-4 (FMO shown in **S4 Fig**). We found that mice infected with IIΔ*gra15* parasites generated a lower frequency of exhausted CD4+ T cells, though the total number of exhausted CD4+ T cells only trended down in IIΔ*gra15* infected mice (**Fig 5A,B**). As NK cells have been shown to contribute to T cell exhaustion in the chronic phase of disease (37), we also quantified NK cell number and frequency, finding a lower frequency of NK in the IIΔ*gra15* infected mice (**Fig 5C, D**). As with the exhausted T cells, the total number of NK cells only trended down in IIΔ*gra15* infected mice (**Fig 5C, D**).

**Figure 5.**
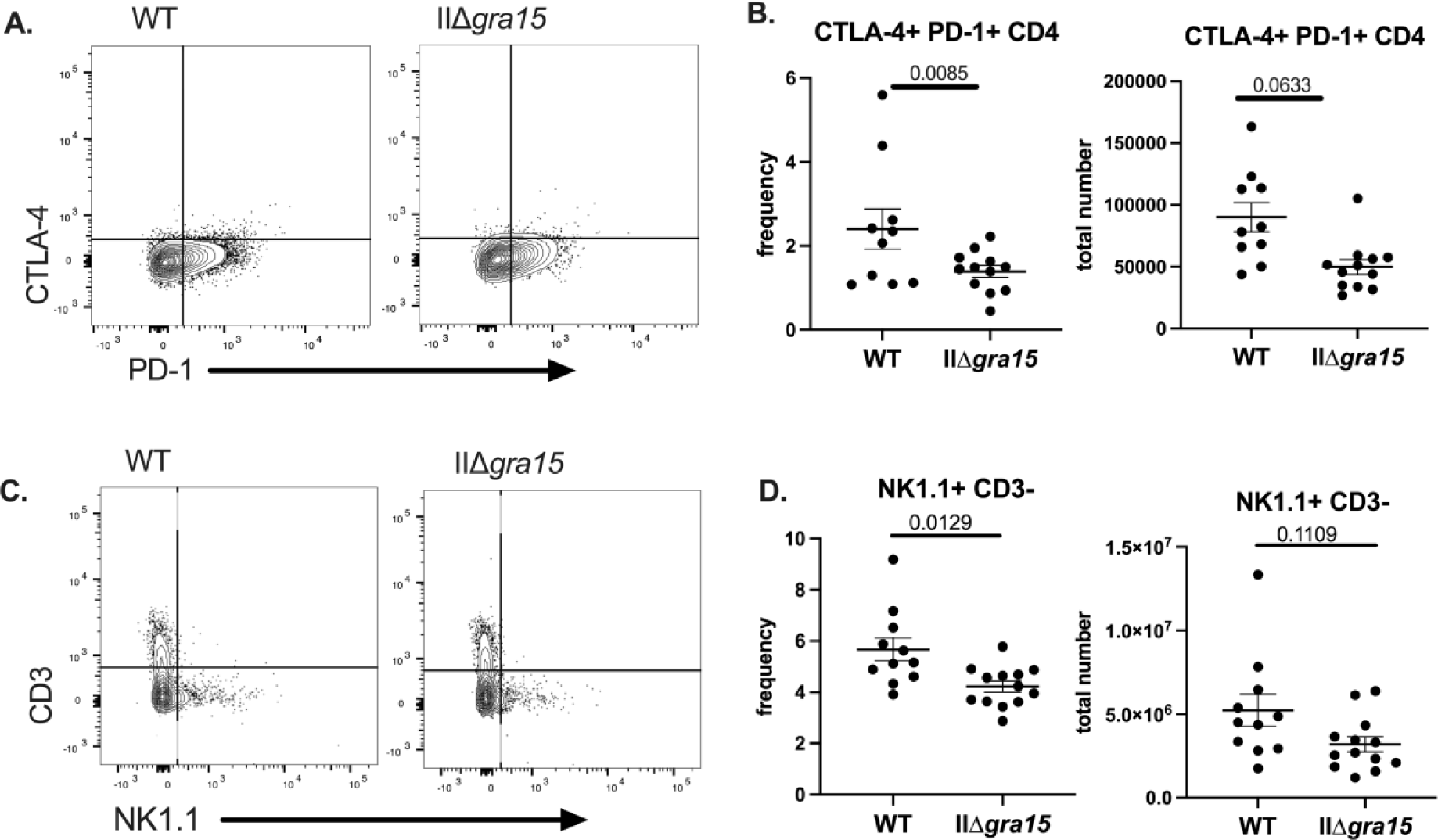
GRA15 may influence the frequency of peripheral “exhausted” T cells and NK cells at 3 weeks post infection. **A,B**. Splenocytes were evaluated for the presence of CTLA-4+/PD-1+ CD4+ T cells **C,D**. Splenocytes were evaluated for the presence of CD3-NK1.1+ cells. Bars, mean ± SEM. N=11-12 mice/infection strain. Data are representative of two independent experiments.

### GRA15 may influence parasite dissemination during acute infection

Given the published data suggesting a difference in parasite burden between WT and IIΔ*gra15* parasites at 5 dpi (14), we were surprised that we did not see a difference in parasite burden in the brain at 3 wpi (**Fig 1A,C**). Therefore, we wondered if the previously reported GRA15-associated phenotypes could only be seen early in infection. To address this question, we inoculated Cre reporter mice intraperitoneally with saline or WT or IIΔ*gra15* parasites and collected peritoneal exudate cells (PECS) and peritoneal fluid. Following the protocol from the previously published report (14), we measured IFN-γ in the peritoneal fluid, finding no difference in IFN-γ levels at 2 dpi (**Fig 6A**). While the prior study used bioluminescent imaging to quantify parasite burden, our parasites were not compatible with such measurements (i.e., our parasites do not express luciferase). Instead, as our parasite strains express a rhoptry::Cre fusion protein and in Cre reporter mice the number of GFP+ cells correlates with the parasite burden (29), we used the number of green fluorescent protein-expressing (GFP+) PECs as an indirect measure of peritoneal parasite burden. Unlike the prior work, at 2 and 5 dpi, we found no difference in the frequency of GFP+ CD45+ PECs between the two groups (**Fig 6B**). Though the GFP+ PEC number were equivalent between WT and IIΔ*gra15* infections at 2 and 5 dpi, Q-PCR for *Toxoplasma* B1 on genomic DNA isolated from liver and spleen at 5 dpi was lower in the IIΔ*gra15* infected mice (**Fig 6C, D**). In summary, unlike previously published data, we did not find a decrease in IFN-γ within the peritoneal cavity at 2 dpi, nor did we find evidence of an increase in the number of IIΔ*gra15* parasites compared to WT parasites at 2 or 5 dpi. On the contrary, if anything, our Q-PCR data suggest the opposite.

**Figure 6.**
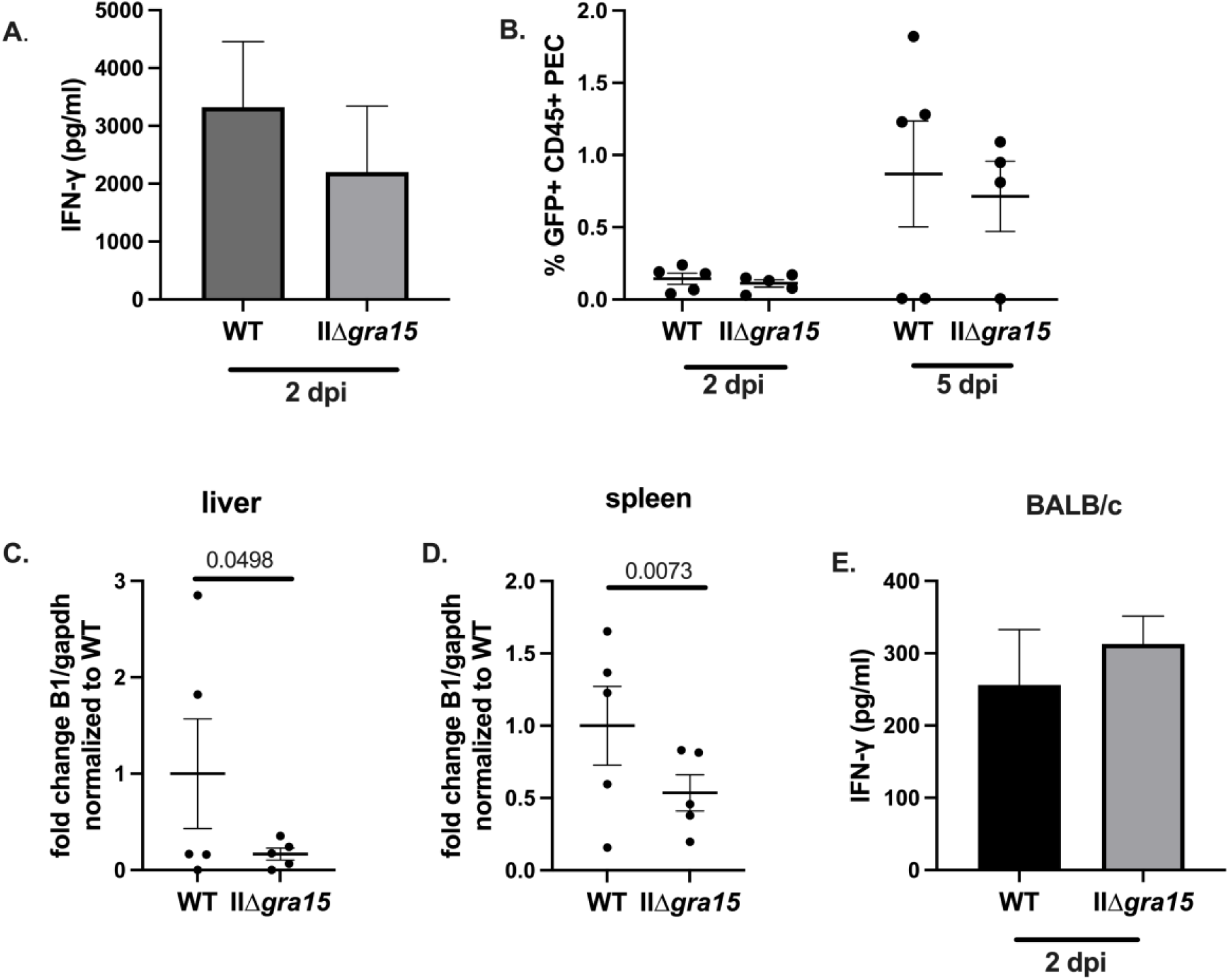
GRA15 may increase parasite dissemination during acute infection. **A-D**. Cre reporter mice were inoculated with 10,000 WT or IIΔ*gra15* parasites. At denoted time points peritoneal lavage was done to isolate peritoneal exudate cells (PECs). At 5 dpi liver and spleen tissue was also collected for B1 analysis. **A.** ELISA of IFN-γ found in the peritoneal cavity at 2 dpi. **B.** Frequency of GFP+ CD45+ cells found within the peritoneal cavity at 2 and 5 dpi. **C.** Q-PCR of parasite genomes on DNA isolated from liver at 5 dpi**. D.** Q-PCR of parasite genomes on DNA isolated from spleen at 5 dpi. **E.** BALB/c mice were intraperitoneally inoculated with 5,000 WT or IIΔ*gra15* parasites. Graph shows levels of IFN-γ detected by ELISA using peritoneal lavage fluid at 2 dpi. Bars, mean ± SEM. N = 4-5 mice/infection strain. **B-D**. Each dot represents a mouse. **A-D**. Data are representative of two independent experiments, 4-5 mice/infection strain/cohort. Statistics: Two-way ANOVA, Fisher’s LSD multiple comparisons test. **E**. Statistics: T-test. N=5 mice per group, one experiment.

Given that our findings were inconsistent with the prior work, we speculated that these discrepancies arose from our using C57BL/6 mice while the prior work used BALB/c mice. We were particularly interested in this possibility because BALB/c mice and C57BL/6 mice are known to generate very different immune responses, with BALB/c mice being predisposed to a Th2 response and C57BL/6 being predisposed to a Th1 response (38–40). To determine if differences in mouse strain explained the discrepancy between our work and the prior work, we inoculated a cohort of BALB/c mice with WT or IIΔ*gra15* parasites and measured IFN-γ levels in the peritoneal cavity at 2 dpi. We found no difference in IFN-γ levels between WT and IIΔ*gra15* infections in BALB/c mice (**Fig 6E**). However, consistent with BALB/c mice being predisposed to generating a Th2 response, the IFN-γ levels in peritoneal fluid of BALB/c mice (**Fig 6E**) was approximately 10-fold lower than IFN-γ levels in the peritoneal fluid of infected C57BL/6 mice (**Fig 6A**).

## Discussion

As GRA15 acutely modulates the secretion of IL-12 by infected macrophages *in vitro* and has been reported to affect parasite growth and local IFN-γ levels very early *in vivo* (15,18,41), here we sought to understand the biological relevance of these changes beyond the earliest days of infection by assessing brain outcomes at 3 wpi. We found that the brain parasite burden, the number of macrophages/microglia and T cells in the brain, and splenic CD4 and CD8 IFN-γ+ T cells did not differ between WT and IIΔ*gra15* strains. We did find several subtle differences in splenocytes from WT and IIΔ*gra15* infected mice (decreased M1-like macrophages and frequency of PD-1+ CTLA-4+ CD4+ T cells and NK cells), but the biological significance of these findings is unclear given the other equivalent outcomes. In summary, the work presented here suggests that despite GRA15’s well documented effects *in vitr*o (14,15,18,32), for the outcomes we measured, GRA15 has little effect on cerebral toxoplasmosis and peripheral immune cell polarization in C57BL/6 mice at 3 wpi.

Our finding that GRA15 does not influence brain parasite burden, at least early in brain infection, is consistent with a prior publication that also used an independently generated II*Δgra15* strain (19). Conversely, a different publication that used strains generated by the lab that originally identified the link between GRA15 and NF-κB found a trend (p>0.05) toward a lower cyst burden at 4 wpi in mice infected with that IIΔ*gra15* strain (14,20). Collectively, these data suggest that GRA15 likely does not influence cyst burden early in brain infection, though variation can be seen with knockouts from different labs.

Though we did not find GRA15-related differences in the brain parasite burden or immune cells in the brain parenchyma, our identification of mice infected with IIΔ*gra15* as having a lower number and frequency of M1-like macrophages (**Fig 2A,B**) is consistent with the *in vitro* data suggesting GRA15 plays a role in polarizing macrophages to an M1-like phenotype (14). However, the rest of the results indicate that this difference in the M1-like compartment is not sufficient to alter parasite abundance in the brain at 3 wpi. Our finding that IIΔ*gra15* infected mice have a lower frequency of “exhausted” CD4+ T cells is novel and interesting, especially when viewed in the context that IIΔ*gra15* infected mice had the same number of IFN-γ producing CD4 and CD8 T cells as WT infected mice (i.e., no IIΔ*gra15* effect on these populations). While several possibilities might explain this discrepancy, one possibility is that these PD-1+ CTLA-4+ CD4 T cells are not exhausted. Recent work suggests the identification of exhausted cells using surface markers only is likely inadequate as PD-1^hi^ cells that also express other inhibitory markers (e.g. TIM-3, CTLA-4) can be highly activated effector cells (i.e., express IFN-γ) that have not yet fully differentiated (42). As our flow panel that included PD-1 and CTLA-4 did not include IFN-γ, we cannot determine if these cells were truly exhausted or maintain effector function. Future studies will be required to definitively determine the status of these cells.

The major limitation of this study, and every study that has examined type II *gra15* knockout strains in mice (14,18–20), is the lack of an appropriate complemented strain in which GRA15 has been ectopically expressed at the same level as in wild-type parasites. While a complemented strain is not necessary for negative results (i.e., no difference between the wild-type and KO strain), complemented strains are important for phenotypes that differ between wild-type and KO strains or between studies of independently generated KOs. For example, as noted above, we found differences between mice infected with the wild-type and IIΔ*gra15* strains in the M1-like, “exhausted” CD4 T cell, and NK cell populations. Similarly, unlike prior work, we did not find a decrease in peritoneal supernatant IFN-γ at 2 dpi in C57Bl/6 or BALB/c mice infected with IIΔ*gra15* parasites compared to mice infected with WT parasites (**Fig 6A, E**). While several possibilities might explain these differences, appropriate complemented strains would help distinguish between GRA15 driven effects and effects driven by idiosyncratic differences of individual knockout strains.

Why does GRA15—which has clear and consistent phenotypes *in vitro* (e.g., NF-κB activation, IL-1β production)—have such a limited phenotype in mice? This discrepancy may relate to *Toxoplasma* having evolved to survive across a range of intermediate hosts, leading to redundancies in mechanisms to manipulate host signaling pathways. For example, *Toxoplasma* proteins GRA83, GRA24, profilin, GRA7, and GRA15 are all linked to IL-12 production from infected murine DCs and macrophages and are initiated through different ligand-receptor interactions (6,14,15,18,43–45). Therefore, these redundancies in how parasites trigger IL-12 production in mice may compensate when parasites lack GRA15. On the other hand, species that lack the receptors that murine cells use to detect *Toxoplasma* protein (e.g., humans lack TLR11/12 which recognize the *Toxoplasma* protein profilin) may have a stronger dependency on GRA15 signaling to generate IL-12 during infection. Thus, while GRA15 may not play an essential role in mice up to 3 wpi, in a different host, GRA15 may be the difference between type II parasite survival and clearance.

## Acknowledgements

We would like to thank the Koshy lab, both past and current members, for continuous discussion, fun, and support. We thank Melissa Lodoen for careful review of and commentary on the manuscript. We also thank the Department of Neuroscience at the University of Arizona for the use of the Imaris software. We thank UBRP student Sakthi Kumar for her help making the IIΔ*gra15* parasite line. We thank John Boothroyd for the gift of mouse anti-Sag1 antibody. We also thank the University of Arizona Cancer Center Flow Cytometry Core (supported by the National Cancer Institute of the National Institutes of Health under award number P30 CA023074) and the UA Equipment Enhancement Fund for Improving Health (Echo Revolve) for enabling parts of this research.

