## Supplementary Figures for "*Toxoplasma* type II effector GRA15 has limited influence *in vivo*"

**Supplementary Figure 1. Evaluation of IIΔ*gra15* and IIΔ*gra15*:GRA15 parasites.**

**A.** Representative image of immunofluorescence assay of human foreskin fibroblast monolayers infected with the listed parasite strains. Twenty-four hours post infection (hpi), cultures were fixed and stained with Hoechst (nuclear stain, blue), anti-NF-κB antibodies (green), and anti-SAG1 antibodies (*Toxoplasma* surface antigen, red). Dotted white line surrounds nuclear NF-κB and nuclear stain. **B.** Graph of nuclear NF-κB mean pixel intensity for the listed parasite strains at 24 hpi. **C**. Graph of nuclear NF-κB mean pixel intensity for the listed parasite strains at 1, 3, and 24 hpi. **B,C.** 35-50 cells analyzed/Toxoplasma strain/time point **D.** Quantification of the number of parasites/parasitophorous vacuole for identified strains at 24 hpi. Bars, mean ± SEM. N>100 vacuoles/well, 3 wells/experiment, 3 experiments. * indicates a significant p value (<0.05) compared to WT **E.** Quantification of plaques formed by identified strains at 10 dpi. **F.** Quantification of total plaque area. **E,F.** Each dot = 1 experiment, N=3 experiments. Bars, mean ± SEM **G.** Representative images of plaque assay. **H.** RNA expression of *gra15* in identified strains. **D.** Two-way ANOVA, uncorrected Fisher's LSD **E,F,H**. One way ANOVA.

**Supplementary Figure 2. Gating scheme of macrophage panel.**

Immune cells were isolated from the spleen and stained for macrophage markers. Single cells were discriminated from doublets by plotting side scatter height (SSC-H) vs side scatter area (SSC-A). Live cells were gated on using live/dead Yellow$^{-}$. CD45+ F4/80+ cells were gated by plotting F4/80 versus CD45. Macrophages were discriminated by plotting the CD45+ F4/80+ population by CD11b versus CD11c. The CD11b$^{hi}$ CD11c$^{lo/int}$ cells were gated on CD80+ CD86+ (M1-like macrophages) or F4/80+ CD206+ (M2 macrophages). Gating scheme shows example of infected sample, saline sample, and the FMO for each final gate.

**Supplementary Figure 3. Gating scheme for CD4 and CD8 T cells**. Splenocytes were gated on singlets, live cells, CD3+, and CD4 versus CD8. CD8 populations were then gated on CD8+/IFN-γ+. CD4 populations were gated on CD4/IFN-γ, CD4/T-bet+, CD4/Gata-3+, or CD4/Foxp3+. Gating scheme shows example of infected sample, saline sample, and the FMO for each final gate.

**Supplementary Figure 4. FMO for denoted T cell gates. A.** FMO for CTLA-4+ and PD-1+ populations. Gates preceding CTLA-4 and PD-1 for CD4 T cells denoted in **Supplementary Figure 3**. **B.** FMO for CD3 and NK1.1 . Cells pre-gated on singlets, live cells.

**Supplementary Figure 5. Frequency and total number of CTLA-4+ PD-1+ CD8 T cells.** Splenocytes were evaluated for the presence of CTLA-4+/PD-1+ CD8+ T cells at 3 wpi.

**Supplementary Figure 6. At 5 dpi, IIΔ*gra15* infected mice have fewer parasite genomes in the liver and spleen compared to WT infected mice. A,B.** Q-PCR of parasite genomes on DNA isolated from liver and spleen from the second cohort of data used for this analysis. The first cohort is depicted in Fig 6 C,D. **A.** One mouse (red dot) was removed from the WT group as it was identified as an outlier by the ROUT Outlier test. Normalization and statistics were run without this data point.

45 **Supplementary Figure 1**

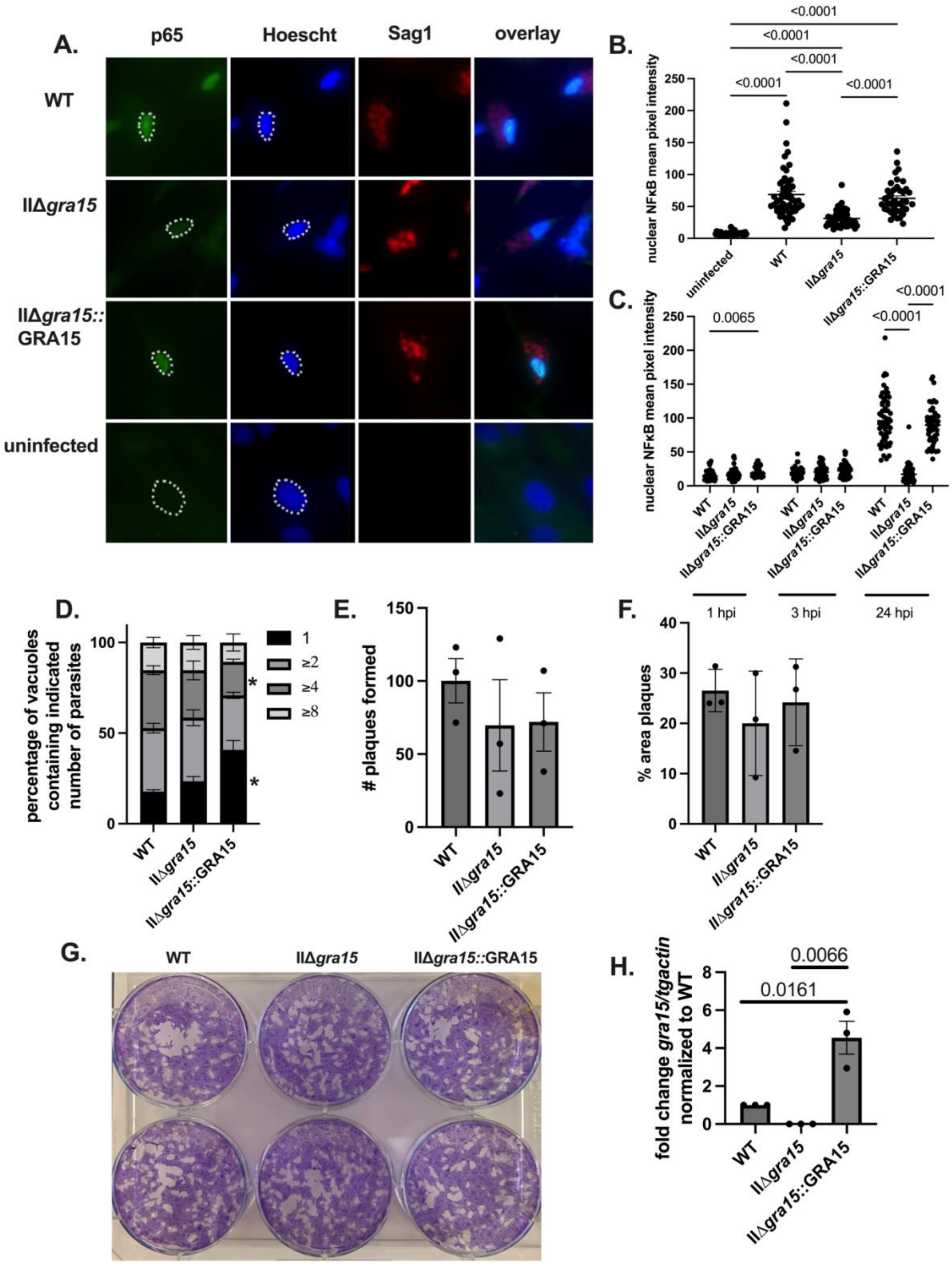


46

47

**Supplementary Figure 2**

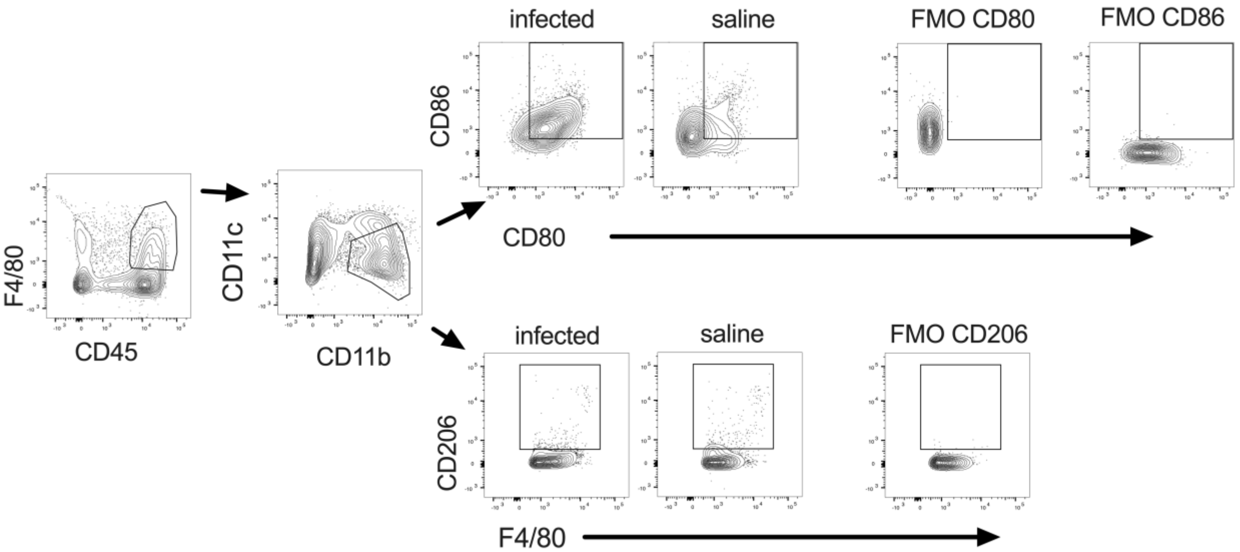

### Supplementary Figure 3

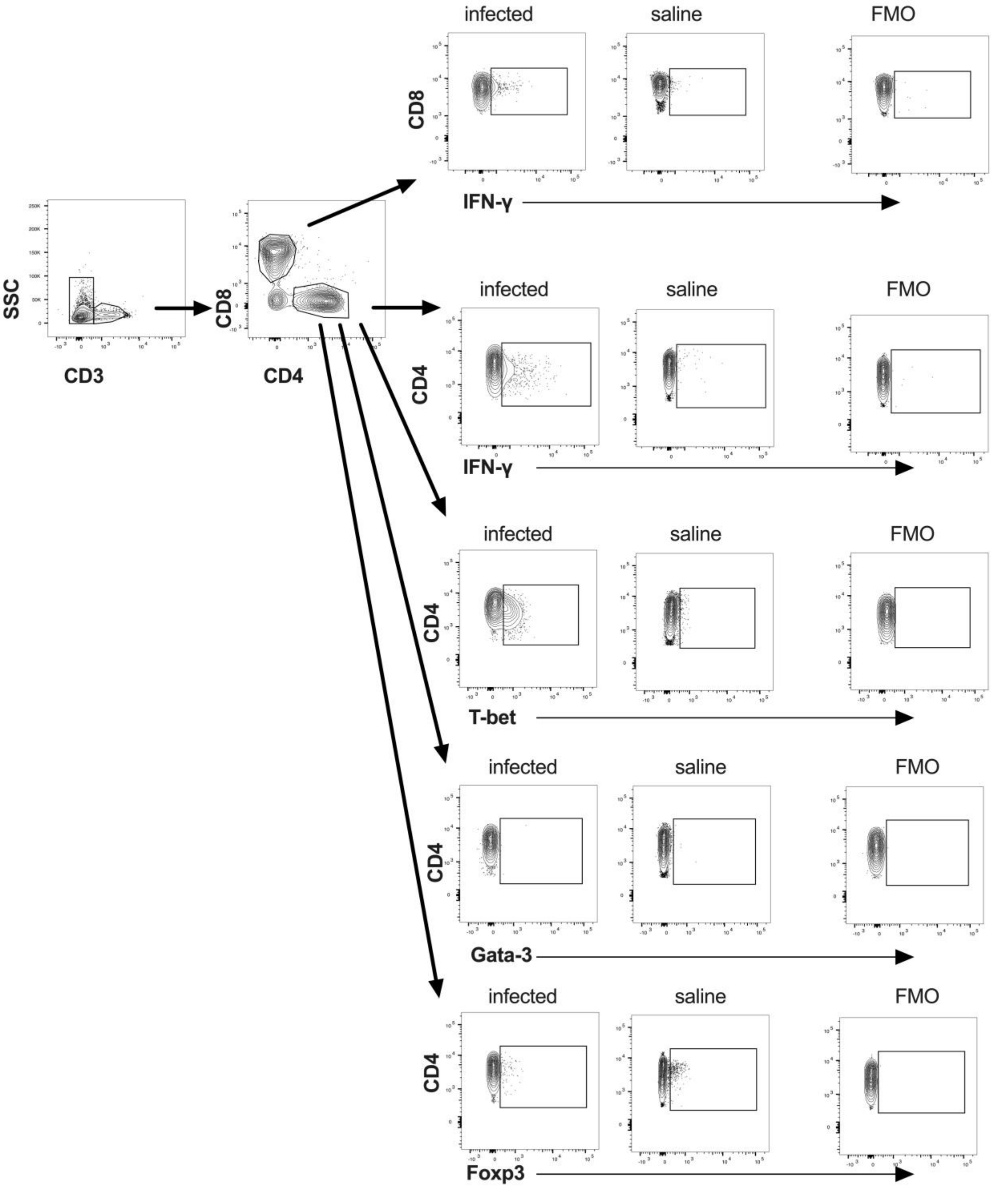

**Supplementary Figure 4**

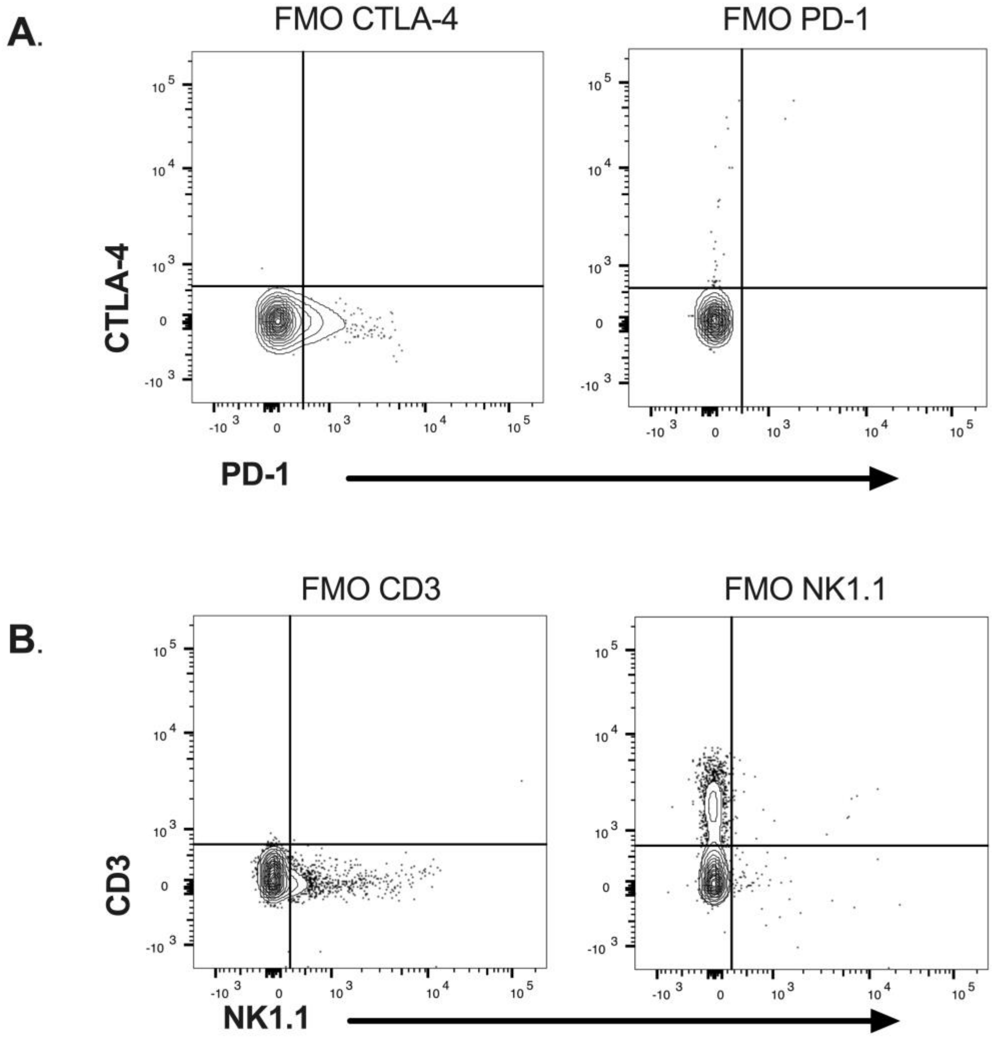

**Supplementary Figure 5**

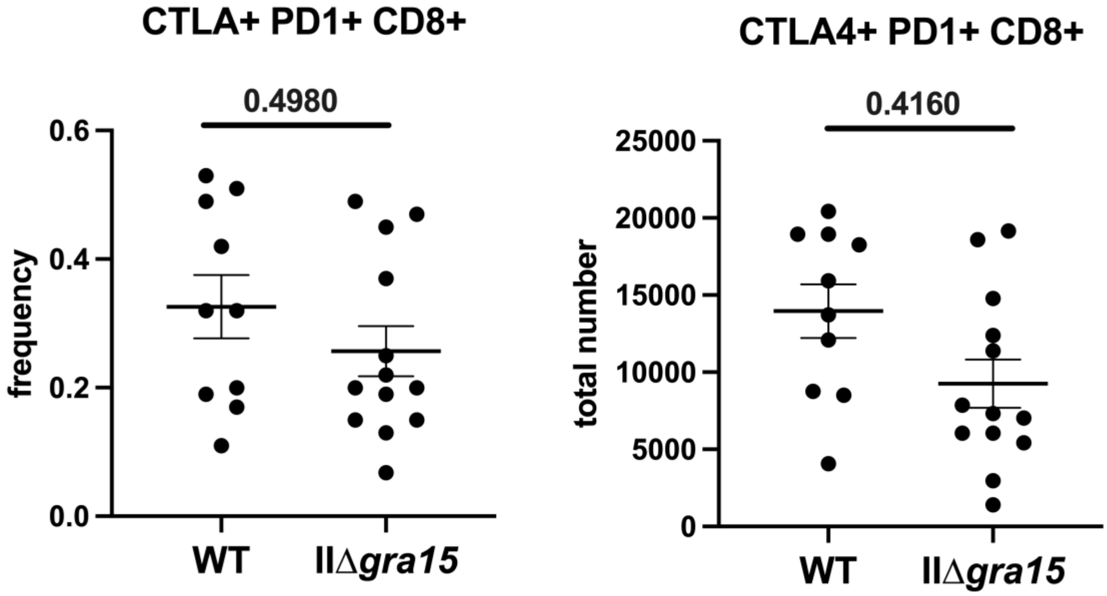

**Supplementary Figure 6**

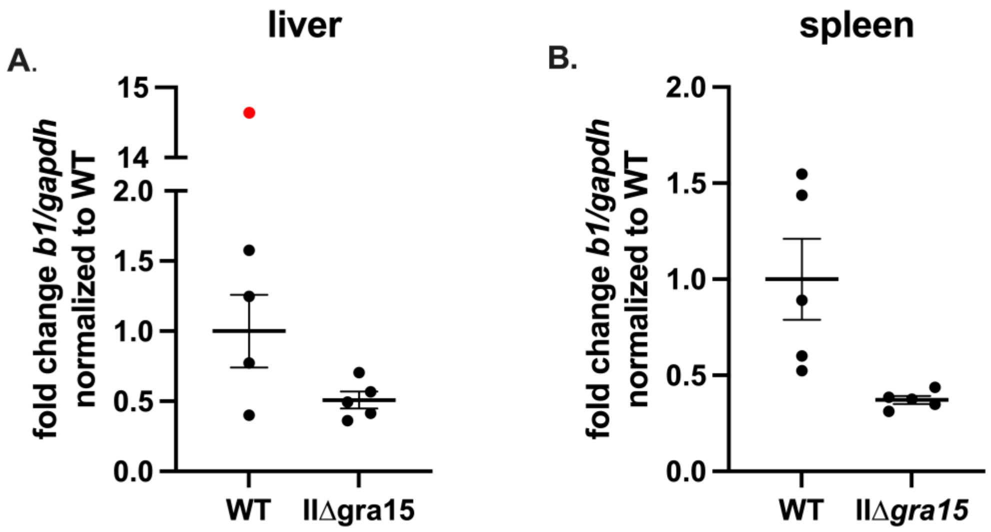
